# Structure of ATP synthase under strain during catalysis

**DOI:** 10.1101/2022.01.24.477618

**Authors:** Hui Guo, John L. Rubinstein

## Abstract

ATP synthases are macromolecular machines consisting of an ATP-hydrolysis-driven F_1_ motor and a proton-translocation-driven F_O_ motor. The F_1_ and F_O_ motors oppose each other’s action on a shared rotor subcomplex and are held stationary relative to each other by a peripheral stalk. Structures of resting mitochondrial ATP synthases revealed a left-handed curvature of the peripheral stalk even though rotation of the rotor, driven by either ATP hydrolysis in F_1_ or proton translocation through F_O_, would apply a right-handed bending force to the stalk. We used cryoEM to image yeast mitochondrial ATP synthase under strain during ATP-hydrolysis-driven rotary catalysis, revealing a large deformation of the peripheral stalk. The structures show how the peripheral stalk opposes the bending force and suggests that proton translocation during ATP synthesis causes accumulation of strain in the stalk, which relaxes by driving the relative rotation of the rotor through six sub-steps within F_1_, leading to catalysis.

## Introduction

ATP synthases use a transmembrane electrochemical proton motive force (pmf) to generate adenosine triphosphate (ATP) from adenosine diphosphate (ADP) and inorganic phosphate (Pi). The enzyme complex consists of two molecular motors positioned to oppose each other’s action on a shared rotor subcomplex (Fig. 1A, *left*). The membrane-embedded F_O_ motor is driven by proton translocation across the membrane through two offset half channels (Junge et al., 1997; Vik and Antonio, 1994) while the soluble F_1_ motor is powered by ATP hydrolysis. In *Saccharomyces cerevisiae*, the F_O_ region contains subunits a, e, f, g, i/j, k, 8, part of subunit b, and the c_10_-ring of the rotor (Liu et al., 2015), while the F_1_ region includes a trimer of catalytic subunit αβ pairs and subunits γ, δ, and ε from the rotor (Abrahams et al., 1994). Coupling between F_1_ and F_O_ requires that the two motors are held stationary relative to each other by a peripheral stalk subcomplex (Fig. 1A, *green structure*), which in yeast is formed from subunits b, d, h, and OSCP (the oligomycin sensitivity conferral protein).

**Figure 1.**
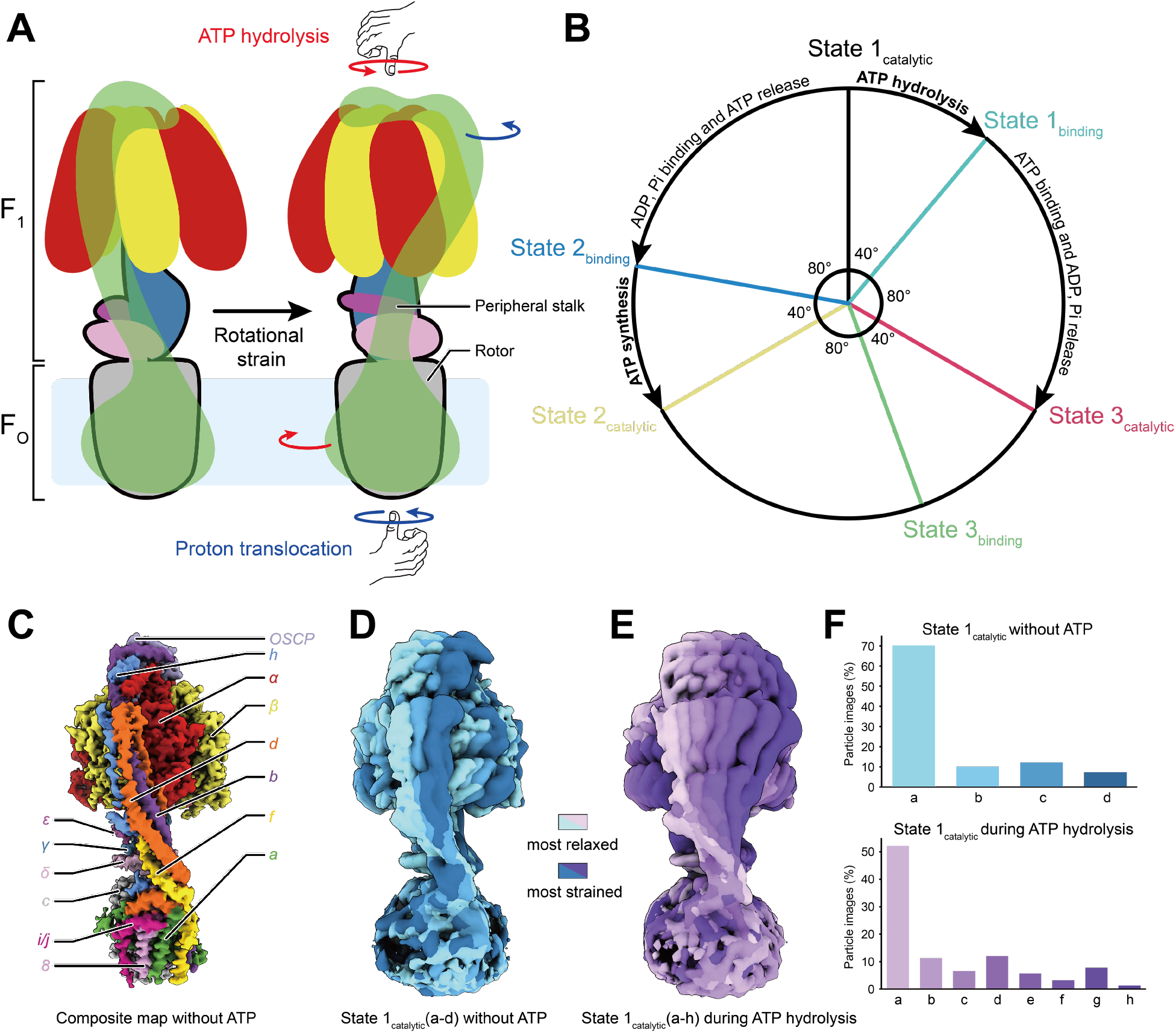
Rotation in ATP synthase. **A,** ATP synthase (*left*) consists of an F_1_ and an F_O_ region with a shared rotor subcomplex (*outlined in black*) and a peripheral stalk (*green*). Rotation driven by proton translocation through the F_O_ region, or the opposite rotation driven by ATP hydrolysis in the F_1_ region, are predicted to induce a right-handed bend of the peripheral stalk (*right*). **B,** ATP hydrolysis or synthesis in the F_1_ region requires three catalytic and three bind dwell conformations. **C**, High-resolution structure of the yeast mitochondrial ATP synthase. **D,** In the absence of free ATP, the peripheral stalk exhibits limited flexibility with a left-handed curvature. **E,** During ATP hydrolysis, ATP synthase can adopt conformations that show a right-handed curvature of the peripheral stalk. **F,** Histograms of the distribution of conformations in the absence of ATP (*top*) and during ATP hydrolysis (*bottom*).

During ATP synthesis, proton translocation through F_O_ at the interface of subunit a and the c-ring causes the γδεc_10_ rotor (Fig. 1A, *outlined in black*) to turn. Rotation of subunit γ within F_1_ leads each αβ pairs to cycle through open, tight, and loose conformations that result in the formation of ATP. Conversely, sequential ATP hydrolysis at each of the three αβ pairs in F_1_ causes the γ subunit to turn in the opposite direction, rotating the proton-carrying c-ring against subunit a in F_O_ and pumping protons across the membrane. Even with the rotor turning at hundreds of revolutions per second (Bilyard et al., 2013; Kobayashi et al., 2020) there is little or no ‘slip’ (Soga et al., 2017) and the H^+^:ATP ratio remains constant. In *S. cerevisiae* this ratio is 10:3 due to the ten proton-carrying c subunits in F_O_ and three catalytic sites in F_1_ (Stock et al., 1999). With this H^+^:ATP ratio, when 10× the free energy of proton translocation 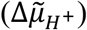 is more negative than 3× the free energy of ATP hydrolysis (Δ*G_ATP_*) the F_O_ motor overpowers the F_1_ motor, forcing it to synthesize ATP. When 3 × Δ*G_ATP_* is more negative than 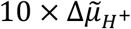, F_1_ overpowers F_O_ and the enzyme hydrolyzes ATP to pump protons.

Synthesis or hydrolysis of each ATP molecule is associated with a ~120° rotation of the γ subunit within F_1_, leading to conformations of the enzyme known as rotational State 1, 2, and 3 (e.g. Zhou et al., 2015). During ATP hydrolysis, which is better studied than ATP synthesis, this ~120° rotation is broken down into a ~40° sub-step as the enzyme transitions from a ‘catalytic dwell’ to a ‘binding dwell’, and an ~80° sub-step as the enzyme transitions to the next catalytic dwell (Yasuda et al., 2001a; Bilyard et al., 2013; Martin et al., 2014; Steel et al., 2015) (Fig. 1B, *clockwise*). ATP hydrolysis likely occurs during the ~40° sub-step while ATP binding likely occurs during the ~80° sub-step (Nishizaka et al., 2004; Adachi et al., 2007; Steel et al., 2015). Consequently, the expected sequence of states for a 360° rotation of the rotor during ATP synthesis is State 1_binding_→State 1_catalytic_ → State 2_binding_ → State 2_catalytic_→State 3_binding_ → State 3_catalytic_ (Fig. 1B, *counter-clockwise*). The mismatch between these six sub-steps in F_1_ and the ten proton-translocation steps in F_O_ suggests that the enzyme cycles between strained and relaxed conformations during catalysis (Cherepanov et al., 1999; Pänke and Rumberg, 1999). Early cryoEM noted that the peripheral stalks of mitochondrial ATP synthases have a left-handed curvature (Lau et al., 2008; Rubinstein et al., 2003) (Fig. 1A, *left*). However, torque applied to the rotor following proton translocation through F_O_ (Fig. 1A, *right, blue arrows*) would tend to rotate the α_3_β_3_ hexamer in the same direction as the torque, inducing a right-handed curvature of the peripheral stalk as it resists the rotation. Similarly, the opposite torque applied to the opposite end of the rotor by ATP hydrolysis in F_1_ (Fig. 1A, *right, red arrows*) would tend to rotate the membrane-embedded region of F_O_ along with the c-ring, also inducing a right-handed curvature of the peripheral stalk as it resists the rotation (Lau et al., 2008). Previously observed structures were obtained in the absence of a pmf or free ATP (Flygaard et al., 2020; Gu et al., 2019; Guo et al., 2019; Hahn et al., 2018, 2016; Lau et al., 2008, p. 20; Mühleip et al., 2021, 2019; Murphy et al., 2019; Pinke et al., 2020; Rubinstein et al., 2003; Sobti et al., 2016; Spikes et al., 2020; Srivastava et al., 2018; Zhou et al., 2015), suggesting that the peripheral stalk may act as a spring that has a left-handed curvature when relaxed but a right-handed curvature under strain during catalysis (Lau et al., 2008).

## Results and discussions

### The peripheral stalk shows pronounced bending under strain during ATP hydrolysis

We purified *S. cerevisiae* ATP synthase with the detergent n-Dodecyl-β-D-Maltopyranoside (DDM), which results in a monomeric preparation of the enzyme (Rubinstein and Walker, 2002; Srivastava et al., 2018), and determined its structure by cryoEM (Fig. S1 and S2, Supplementary Table 1 and 2). A high-resolution map of the intact complex was generated by combining multiple maps from focused refinements (Fig. 1C and S1C). In this map, the peripheral stalk shows the left-handed curvature seen previously. Three-dimensional (3D) classification allowed particle images to be separated into six rotor positions, corresponding to the catalytic and binding dwells for each of the three main rotational states. These conformations resemble recent catalytic and binding dwell structures for an isolated bacterial F_1_ subcomplex imaged during ATP hydrolysis, where the absence of the peripheral stalk resulted in all catalytic dwell structures being identical and all binding dwell structures being identical (Sobti et al., 2021). For yeast ATP synthase imaged without ATP, the catalytic dwell structures show αβ_tight_ either in the open conformation lacking nucleotide or in a closed conformation with weak nucleotide density, and the binding dwell structures show αβ_tight_ only in an open conformation without nucleotide (Fig. S3). The existence of αβ_tight_ in an open conformation without visible nucleotide density is likely an artefact from loss of ATP during the purification of the enzyme. Further classification of the State 1_catalytic_ conformation resulted in classes distinguished by variability in the position of the peripheral stalk and a slight rotation of the rotor relative to subunit a. These classes were designated as State 1_catalytic_(a) (Fig. 1D, *light blue*) to State 1_catalytic_(d) (Fig. 1D, *dark blue*) in order of increasing straightening of the peripheral stalk (Supplementary Video 1, *‘no ATP’ condition*). As these structures were determined in the absence of free ATP, they likely represent energetically similar conformations that can be reached by thermal fluctuation of the enzyme structure (Murphy et al., 2019; Zhou et al., 2015).

To test the hypothesis that the peripheral stalk of ATP synthase deforms under strain, we next added 10 mM ATP to the preparation and froze cryoEM specimens within 10 s. Analysis of a small dataset of images for this condition collected with a screening electron microscope revealed conformations of the enzyme not seen in the absence of free ATP (Fig. S4). Therefore, a large dataset was collected for the specimen with a high-resolution microscope (Fig. S5). Classification of the resulting dataset yielded maps showing six different F_1_ states, corresponding to the catalytic and binding dwell structures from each of the three main rotational states. Subclassification of these populations separated each catalytic and each binding state into conformations with increasing rotation of the rotor relative to subunit a, and increasingly strained peripheral stalks, labelled as ‘a’, ‘b’, ‘c’, etc. Overall, 27 unique conformations were identified: State 1_binding_(a to d), State 1_catalytic_(a to h), State 2_binding_(a to b), State 2_catalytic_(a to e), State 3_binding_(a to c), and State 3_catalytic_(a to e). Overlaying the eight State 1_catalytic_ (a to h) structures reveals that during ATP hydrolysis the peripheral stalk exhibits a large bending motion, transitioning from a left-handed curvature (Fig. 1E, *light purple*) to the predicted right-handed curvature (Fig. 1E, *dark purple*; Supplementary Video 1, *‘During ATP hydrolysis’ condition*). Without ATP and during ATP hydrolysis, the left-handed curvature of the peripheral stalk remains the most highly populated conformation of the enzyme (Fig. 1F).

### The flexible peripheral stalk accommodates rigid rotation of the rotor during ATP hydrolysis

To facilitate comparison of the ATP synthase conformations that occur during ATP hydrolysis, backbone models of the protein structure were fit flexibly into each of the 27 maps (Fig. 2A). Remarkably, the α_3_β_3_γδεc_10_ models from all 18 catalytic dwell conformations could be overlaid with high-fidelity (Fig. 2B, *left*), as could the nine α_3_β_3_γδεc_10_ models from binding dwell conformations (Fig. 2B, *right*), with some limited flexibility at the interface between F_1_ and the c-ring. This observation shows that, other than being in a catalytic or binding dwell conformation, the differences between the structures are mostly due to deformation of the peripheral stalk subunits and the rotation of the c-ring relative to subunit a in F_O_. Comparison of the eight State 1_catalytic_ models shows that the α_3_β_3_γδεc_10_ rotor can turn ~80° against subunit a in F_O_, or more than one fifth of a complete revolution, before transition to the next binding dwell conformation (Fig. 2C). Bending of the peripheral stalk and not the central rotor of the complex supports suggestions that the peripheral stalk is the most compliant part of the enzyme and stores energy during rotary catalysis (Guo et al., 2019; Hahn et al., 2018; Murphy et al., 2019; Sobti et al., 2016; Sorgen et al., 1999, 1998; Zhou et al., 2015).

**Figure 2.**
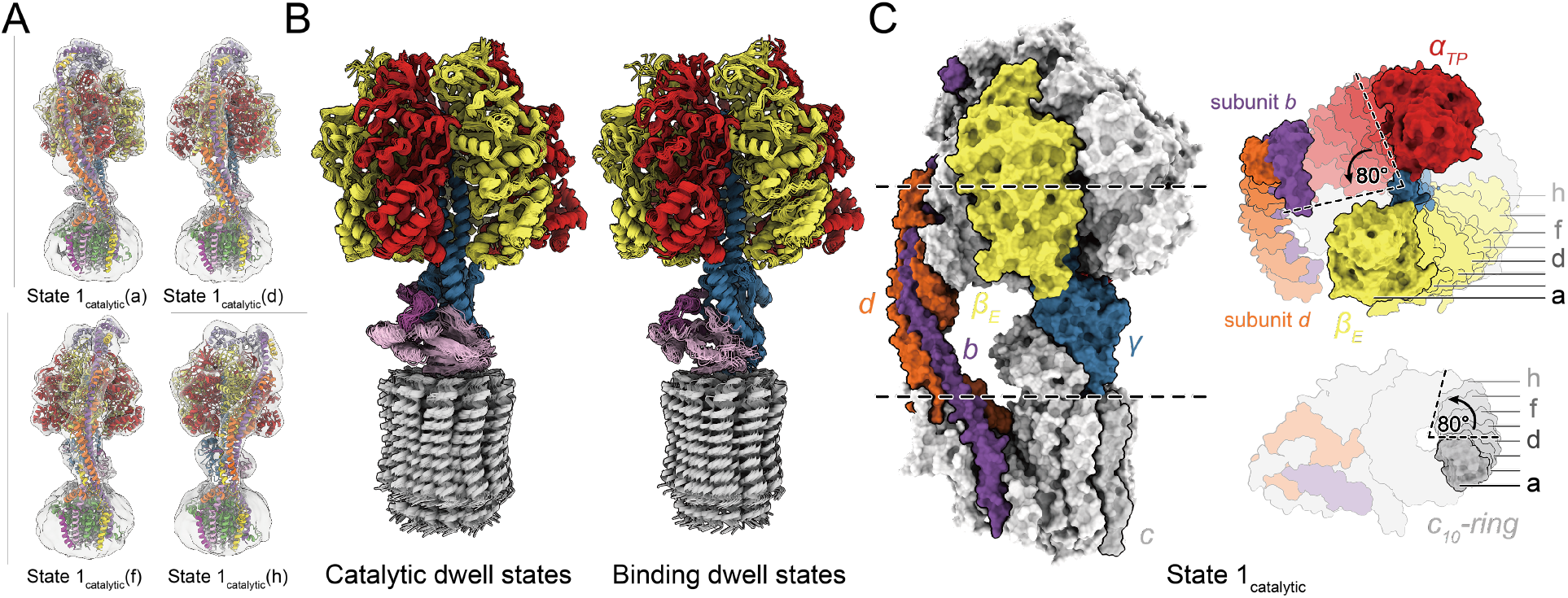
Atomic models of ATP synthase under strain during ATP hydrolysis. **A,** Atomic models of the enzyme were generated by flexible fitting into maps of the strained ATP synthase conformations. **B,** Overlay of fitted models from catalytic dwell (*left*) and binding dwell (*right*) structures show limited conformational changes in the α_3_β_3_γδεc_10_ subcomplex. **C,** Overlay of models from catalytic dwell conformations show pronounced bending of the peripheral stalk while the enzyme is under strain during ATP hydrolysis.

### The peripheral stalk bends by deformation of subunits d, f, and h

The peripheral stalk of yeast ATP synthase contains subunits b, d, h, and OSCP (Fig. 3A). Although atomic models for subunits b, d, and OSCP have been constructed from previous cryoEM of ATP synthase (Srivastava et al., 2018), model quality for the 92-residue subunit h in earlier structures was low due to flexibility in both the peripheral stalk overall and subunit h specifically. Focused refinement of the peripheral stalk in the current structure resulted in continuous density for most of subunit h, allowing for construction of an atomic model for residues 1 to 62 based on predictions from AlphaFold (Jumper et al., 2021) (Fig. 3A, *blue*; Fig. S6). Interestingly, despite density immediately C-terminal of His62 in subunit h appearing disordered, an additional density that interacts with subunits a, d, f, and 8 indicates that the C terminus of the protein reaches the membrane surface, as suggested previously (Rubinstein et al., 2005) (Fig. 3A, *dashed box*). Therefore, subunit h spans the entire distance from F_1_ to F_O_, a role usually attributed only to subunit b, and different from subunit F_6_, the shorter mammalian homologue of subunit h (Spikes et al., 2020).

**Figure 3.**
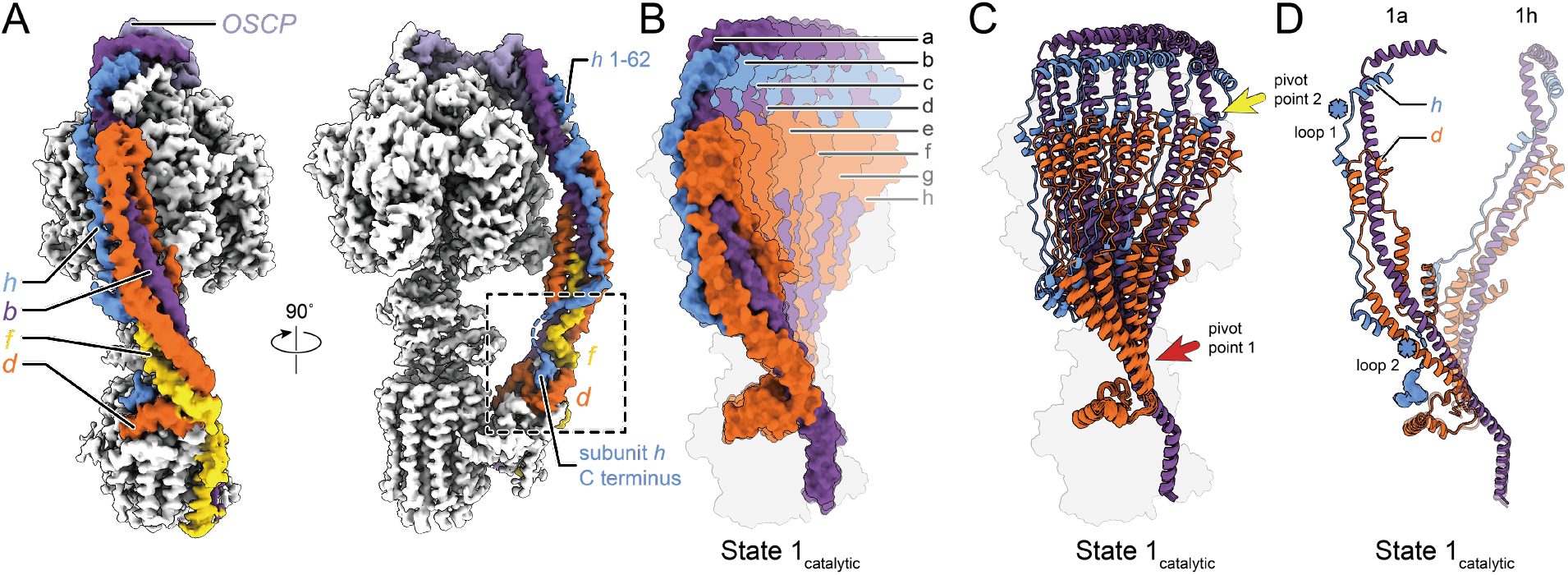
Deformation of the peripheral stalk of ATP synthase while under strain during ATP hydrolysis. **A,** Structure of the peripheral stalk shows that subunit h bridges F_1_ and F_O_. **B,** Overlay of the eight State 1_catalytic_ conformations shows a large deformation of the peripheral stalk. **C,** The peripheral stalk bends at two pivot points near F_1_ (*yellow arrow*) and near F_O_ (*red arrow*). **D,** Unstructured regions in subunit h (*blue asterisks*) allow it to withstand the bending of subunit d.

Aligning the eight structures corresponding to the State 1_catalytic_ by their F_O_ regions reveals that the dramatic bending of the peripheral stalk is facilitated mainly by deformation of subunits b, d, and h (Fig. 3B; Supplementary Video 1, *‘During ATP hydrolysis’ condition*). In conformations that show only slight bending of the peripheral stalk, such as State 1_catalytic_(b to d), deviation from the relaxed State 1_catalytic_(a) conformation is mediated primarily by a pivot point in subunits b and d close to the membrane surface (Fig. 3C, *red arrow*). In the more strained conformations like State 1_catalytic_(h), a second pivot point in subunit b at the top of subunit d is apparent (Fig. 3C, *yellow arrow*). The two pivot points are located at either end of subunit d, indicating that subunit d controls where the peripheral stalk bends and likely acts to oppose the bending force, inducing the left-handed curvature of the peripheral stalk when it is not under strain. The structure of subunit d, with an α-helical hairpin that allows it to push against subunit b, is ideally optimized for its role of applying a force that attempts to restore the relaxed conformation of the peripheral stalk during ATP hydrolysis or synthesis (Fig. 3D; Supplementary Video 1, *orange subunit*). Subunit h contains two disordered regions close to the two pivot points defined by subunit d, which allows it to withstand the large conformational changes that occurs around the pivot points (Fig. 3D, *blue asterisks*). In contrast with the spring-like peripheral stalk seen here for the yeast ATP synthase, the unusually large peripheral stalk of algal ATP synthase from *Polytomella* sp., although imaged in the absence of substrate, appears mostly rigid, with the OSCP subunit that connects the catalytic domain to the rest of the peripheral stalk showing the most flexibility (Murphy et al., 2019).

### Overall rotation cycle of yeast ATP synthase

Despite the presence of a high concentration of ATP in the buffer used for freezing specimens during ATP hydrolysis, State 1_catalytic_(a), the least strained of the State 1_catalytic_ conformations, appears to have MgADP bound in its αβ_tight_ site (Fig. 4A, *left*). In contrast, refinement of the F_1_ region with particle images combined from State 1_catalytic_(e to h), the four most strained of the State 1_catalytic_ conformations, resulted in a structure similar to State 1_catalytic_(a) but with what appears to be MgATP bound to αβ_tight_ (Fig. 4A, *right*). In the presence of free ATP, ATP hydrolysis occurs at the αβ_tight_ site and MgADP within the site is expected to inhibit this hydrolytic activity. Therefore, the presence of MgADP in αβ_tight_ of the non-strained conformation suggests that many of the complexes in this conformation are in the well-known MgADP inhibited state (Abrahams et al., 1994; Bowler et al., 2007). Similarly inactive complexes have been detected previously even in the presence of free ATP (Hirono-Hara et al., 2001; Noji et al., 1997). In contrast, the structures that show the more strained peripheral stalks appear to be calculated from images of active enzyme particles. Density for the binding dwell conformations suggests that they contain MgADP with Pi in the αβ_tight_ site (Fig. S7A), as was seen in the bacterial F_1_ region during ATP hydrolysis (Sobti et al., 2021).

**Figure 4.**
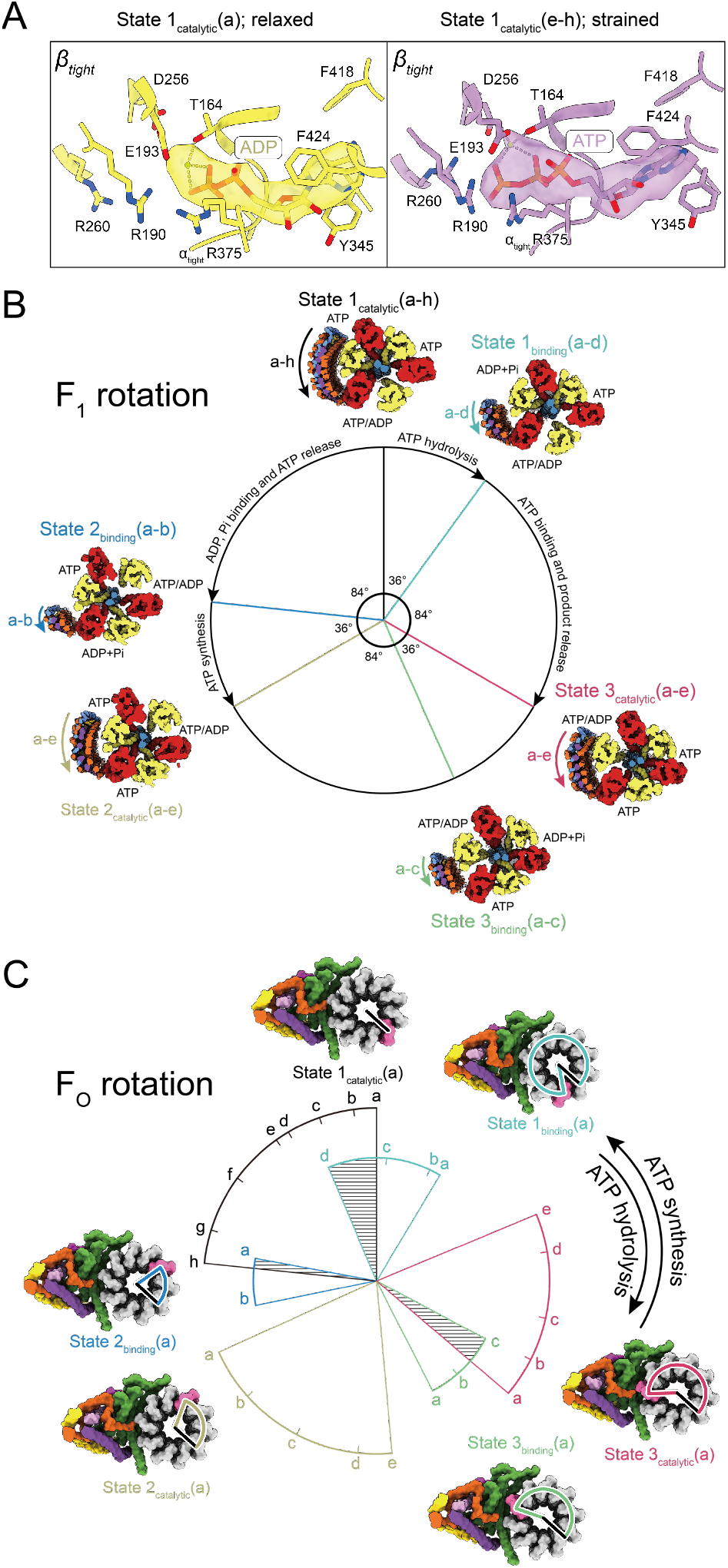
Sequence of conformations in the ATP hydrolysis and ATP synthesis cycles. **A,** The unstrained conformation of State 1_catalytic_ shows density consistent with MgADP in the αβ_tight_ catalytic pair (*left*), suggesting an ADP-inhibited state, while the strained conformations show density consistent with MgATP (*right*) suggesting an active state. **B,** Measurement of the rotation of the rotor subunit g within the F_1_ region shows 36° and 84° sub-steps between catalytic and binding dwell conformations. **C,** Measurement of the rotation of the c-ring relative to subunit a in F_O_. Within each binding or catalytic dwell conformation the peripheral stalk becomes increasingly strained as the c-ring rotates in the ATP synthesis direction and increasingly relaxed as the c-ring rotates in the ATP hydrolysis direction. Transition between catalytic and binding dwell conformations would require back-stepping of the c-ring (shaded areas) if every conformation occurred during the rotary cycle.

To place the 27 conformations of ATP synthase observed during ATP hydrolysis into a rotational sequence, the positions of subunit γ relative to α_3_β_3_ in F_1_ (Fig. 4B) and of the c-ring relative to subunit a in F_O_ (Fig. 4C) were measured and plotted on circles that represent a 360° rotation. As described above, the α_3_β_3_γδεc_10_ subcomplex is found in three catalytic dwell conformations and three binding dwell conformations, resulting in only six unique positions of subunit γ relative to α_3_β_3_ in F_1_ (Fig. 2B). Consistent with the isolated bacterial F_1_ region (Sobti et al., 2021; Yasuda et al., 2001b), ATP hydrolysis in αβ_tight_ of the yeast catalytic dwell conformation appears to induce a slightly more open conformation of the αβ pair and a ~36° rotation of the rotor (Fig. S7B), leaving the enzyme in a binding dwell. MgADP and Pi are then released from the αβ_tight_ site and ATP binding to the αβ_open_ site drives an ~84° rotation of the rotor to the next catalytic dwell conformation. Repetition of this process two more times completes the 360° rotation cycle for ATP hydrolysis (Fig. 4B, *clockwise arrows*), while for ATP synthesis the reverse reaction is driven by rotation of the rotor in the opposite direction (Fig. 4B, *counter-clockwise arrows*).

In contrast to the six unique positions of subunit γ relative to α_3_β_3_ in F_1_, there are 27 unique positions of the c-ring relative to subunit a in F_O_. Plotting the angle of the c-ring relative to subunit a in F_O_ produces a series of arcs that show the range of rotation of the ring within each catalytic or binding dwell state (Fig. 4C, *black, blue, yellow, green, red, and cyan arcs*). These arcs reveal that as the c-ring rotates in the ATP hydrolysis direction, each state exhibits a decreasing strain on the peripheral stalk (Fig. 4C, *clockwise arrow*). For example, for State 1_catalytic_ (Fig. 4C, *black arc*), rotation of the c-ring in the ATP hydrolysis direction occurs during the transition from State 1_catalytic_(h) → State 1_catalytic_(a). As ATP hydrolysis in a catalytic αβ_tight_ site causes the transition from a catalytic dwell to a binding dwell, the order of states indicates that the power stroke of ATP hydrolysis forces the peripheral stalk into a more strained conformation (e.g. State 1_catalytic_(a) → State 1_binding_(d)). This strain subsequently relaxes as the c-ring continues to turn in the ATP hydrolysis direction (e.g. State 1_binding_(d) → State 1_binding_(a)). Conversely, rotation of the c-ring in the direction driven by proton translocation during ATP synthesis (Fig. 4C, *counter-clockwise arrow*) leads to increasing strain on the peripheral stalk (e.g. State 1_binding_(a) → State 1_binding_(d)), which relaxes as ATP is formed in the catalytic site and the enzyme transitions from a binding dwell conformation to a catalytic dwell conformation (e.g. State 1_binding_(d) → State 1_catalytic_(a)).

Notably, the most strained conformation of some of the states show less rotation of the c-ring in the ATP hydrolysis direction than less strained conformations of the preceding state (Fig. 4C*, shaded areas*). For example, the transition from State 1_catalytic_(a) to State 1_binding_(d) during ATP hydrolysis would involve the c-ring rotating 23° in the ATP synthesis direction. The same apparent ‘backstepping’ can be seen at the transition from State 3_catalytic_ → State 3_binding_, and State 2_binding_ → State 1_catalytic_. This backstepping of the c-ring would bend the peripheral stalk in the opposite direction of the applied force and is physically unlikely. Therefore, the unstrained conformations appear to show inactive complexes that are not part of the rotary sequence during substrate turnover. By extension, these data suggest that during rotary catalysis the peripheral stalk becomes strained and does not relax fully until catalysis stops. Construction of a movie showing rotation in the hydrolysis direction based on the most strained conformation of the enzyme illustrates the amount of deformation that can occur during ATP hydrolysis (Supplementary Video 2, *‘ATP hydrolysis’* cycle). Similarly, a video can be constructed showing rotation in the ATP synthesis direction based on the most strained conformations (Supplementary Video 2, *‘ATP synthesis’* cycle). Together, these data illustrate how in active ATP synthase the peripheral stalk can serve as a buffer that deforms under strain. In the fully active enzyme, the peripheral stalk likely remains deformed as the enzyme runs, with the degree of bending dependant on the rate of turnover, and with the enzyme only becoming fully relaxed in the absence of ATP or a proton motive force.

## Methods

### Yeast growth and ATP synthase purification

Yeast was grown and ATP synthase was purified as described previously with minor modifications (Lau et al., 2008; Rubinstein et al., 2005). Briefly, yeast strain USY006 containing a 6×His tag at the N terminus of the β subunits was grown in YPGD media (1% [w/v] yeast extract, 2% [w/v] peptone, 3% [v/v] glycerol, 0.2% [w/v] glucose) with a 11 L fermenter (New Brunswick Scientific) for ~48 hours at 30 °C until saturation. All purification steps were performed at 4 °C. Yeast cell walls were broken with bead beating, and cell debris was removed by centrifugation at 5,000 ×g for 30 min. Mitochondria were collected by centrifugation at 25,000 ×g for 30 min, before being washed with phosphate buffer (50 mM sodium phosphate pH 9.0, 5 mM 6-aminocaproic acid, 5 mM benzamidine, 1 mM PMSF) for 30 min. Washed mitochondria were collected by centrifugation at 184,000 ×g for 30 min, before being resuspended in buffer (50 mM Tris-HCl pH 7.4, 10% [v/v] glycerol, 1% [w/w] DDM [Anatrace], 5 mM 6-aminocaproic acid, 5 mM benzamidine, 1 mM PMSF) and solubilized with gentle shaking for one hour. Insoluble material was removed by centrifugation at 184,000 ×g for 30 min, and supernatant containing solubilized protein was supplemented with 40 mM imidazole and 300 mM NaCl before being loaded onto a 5 mL HisTrap column (Cytiva) equilibrated with HisTrap buffer (50 mM Tris-HCl pH 7.4, 10% [v/v] glycerol, 0.05% [w/w], 40 mM imidazole, 300 mM NaCl, 5 mM 6-aminocaproic acid, 5 mM benzamidine, 1 mM PMSF) and washed with HisTrap buffer. ATP synthase was eluted with HisTrap buffer containing 300 mM imidazole and was loaded onto a Superose 6 Increase column (Cytiva) equilibrated with buffer (20 mM Tris-HCl pH 7.4, 10% [v/v] glycerol, 0.05% [w/w] DDM, 100 mM NaCl, 5 mM MgCl_2_). Fractions containing ATP synthase were pooled, and the protein was concentrated to ~15 mg/ml prior to cryoEM grid freezing or storage at −80 °C.

### CryoEM specimen preparation

Glycerol in the ATP synthase preparation was removed with a Zeba Spin desalting column (Thermo Fisher Scientific [TFS]) before freezing cryoEM specimens. Holey gold films with ~2 μm holes were nanofabricated as described previously (Marr et al., 2014) on 300 mesh Maxtaform copper-rhodium grids (Electron Microscopy Sciences). Specimens with 10 mM ATP were prepared by first applying 0.4 μL of 50 mM ATP in buffer (70 mM Tris-HCl pH 7.4, 0.05% [w/w] DDM, 100 mM NaCl, 55 mM MgCl_2_) onto a grid that had been glow-discharged in air for 2 min. Freshly prepared ATP synthase (1.6 μL) was mixed quickly with the ATP solution on the grid before blotting for 1 s in an EM GP2 grid freezing device (Leica) at 4 °C and 100% humidity and plunge frozen in liquid ethane. Specimens without ATP were prepared the same way except that the mixing step was omitted.

### CryoEM data collection

Preliminary cryoEM data was collected with FEI Tecnai F20 electron microscope operated and 200 kV and equipped with a Gatan K2 Summit camera. Images with this microscope were acquired as movies with 30 fractions at 5 e/pixel/s and a calibrated pixel size of 1.45 Å/pixel. CryoEM movies for high-resolution analysis were collected with a Titan Krios G3 microscope operated at 300 kV and equipped with a Falcon 4 camera (TFS). Automated data collection was performed with *EPU*. For the dataset including ATP, 10,037 movies, each consisting of 30 fractions, were collected at a nominal magnification of 59,000 ×, corresponding to a calibrated pixel size of 1.348 Å. The exposure rate and the total exposure of the specimen were 6.1 e^-^ /pixel/s and ~40 e^-^/Å^2^, respectively. For the ATP-free dataset, 8,817 30-fraction movies were collected at a nominal magnification of 75,000×, corresponding to a calibrated pixel size of 1.046 Å. The exposure rate and the total exposure for this specimen were 4.2 e/pixel/second and ~39 e/Å^2^, respectively.

### Image analysis

Data collection was monitored with *cryoSPARC Live* (Punjani et al., 2017) to screen and select high-quality micrographs. All other image analysis steps were performed with *cryoSPARC v2* except where mentioned. Movie fractions were aligned with patch-based motion correction and contrast transfer function (CTF) parameters were estimated with patch-based CTF estimation. After removing movies with undesirable motion or CTF fit, 7,474 and 4,059 movies from the dataset including ATP and ATP-free dataset were selected for further processing, respectively. Movie fractions were aligned with *MotionCor2* (Zheng et al., 2017) with a 7×7 grid and averaged micrographs from the aligned movies were subjected to patch-based CTF estimation. For the ATP-free dataset, particle selection was perform with Topaz (Bepler et al., 2019). For the dataset including ATP, templates for particle selection were generated from 2D classification of manually selected particle images. After particle selection, 2,534,488 particle images were extracted for the dataset with ATP and 442,025 particle images were extracted for the ATP-free dataset. Low quality particle images were removed with two rounds of 2D classifications, yielding 1,109,677 and 422,765 particle images for the dataset including ATP and the ATP-free dataset, respectively. Further cleaning with ab initio 3D classification and heterogeneous refinement reduced dataset sizes to 915,825 and 379,817 particle images, respectively. The remaining particle images were classified into three classes, corresponding to the three main rotational states of the enzyme, and each class was refined with non-uniform refinement (Punjani et al., 2020). For the dataset including ATP, local refinement was performed with all particle images with a mask including α_3_β_3_γδε from the F_1_ region. CTF parameters of individual particle images were re-estimated with local CTF refinement, and masked refinement was performed again with updated CTF parameters. Image alignment parameters were then converted to *Relion* (Scheres, 2012) .star file format with *pyem* (DOI: 10.5281/zenodo.3576630) and individual particle motion was corrected with Bayesian polishing (Zivanov et al., 2019). For the ATP-free dataset, Bayesian polishing was performed with an intact ATP synthase map reconstructed with all particle images and particle images were down-sampled to a pixel size of 1.308 Å. Motion corrected images were imported back to *cryoSPARC*, refined, and CTF parameters re-estimated. For the dataset including ATP, particle images were initially classified into four classes. Iterative ab initio classification and heterogeneous refinement of each of the four classes yielded 27 unique rotational states of ATP synthase. The 27 structures were named State 1_binding_(a to d), State 1_catalytic_(a to h), State 2_binding_(a to b), State 2_catalytic_(a to e), State 3_binding_(a to c), and State 3_catalytic_(a to e), and had resolutions ranging from 4.4 to 7.8 Å after refinement. Masked local refinement of the F_1_ region of State 1_catalytic_(a) yielded a 3.5 Å resolution map representing the MgADP inhibited state. Local refinement of the F_1_ region of combined State 1_catalytic_(e to h) and State 1_binding_(a to d) yielded two 4.0 Å resolution maps. To better visualize the nucleotide density in maps, density modification (Terwilliger et al., 2020) of locally refined maps of State 1_catalytic_(a), State 1_catalytic_(e to h), and State 1_binding_(a to d) was performed in *Phenix* (Adams et al., 2010). For the ATP-free dataset, a similar 3D classification strategy yielded nine F_1_ states, namely State 1, 2, and 3_catalytic_ with αβ_tight_ in a closed conformation, State 1, 2, and 3_catalytic_ with αβ_tight_ in an open conformation, and State 1, 2, and 3_binding_ with αβ_tight_ in an open conformation (Fig. S1, S3). These states included 56,739, 52,468, 24,879, 31,559, 47,065, 65,651, 19,922, 57,468 and 23,622 particle images, respectively. Local refinement of the F_1_ region with these images yielded maps at 3.4, 3.4, 3.6, 3.5, 3.5, 3.4, 3.7, 3.4, and 3.7 Å resolution, respectively. When the three rotational states are combined, the F_1_ regions of State_catalytic_ with β_tight_ closed, State_catalytic_ with β_tight_ open, and State_catalytic_ with βbinding open reached 3.2, 3.2, 3.3 Å resolution, respectively. Classification of particle images contributing to State 1_catalytic_ from the ATP-free dataset yielded 4 classes with different c-ring positions relative to subunit a, which demonstrates the flexibility of the peripheral stalk in the absence of free ATP (Fig 1D, S1). Maps of these states were calculated from 41,506, 8,810, 10,550, and 1,984 particle images and reached 3.8, 4.4, 4.4, 7.1 Å resolutions after refinement, respectively. A similar classification strategy was employed with the other two catalytic states, and particle images of the most relaxed State 1, 2, 3_catalytic_(a) (191,939 particle images) were used to calculate locally-refined maps of OSCP with its contact site on F_1_, the remainder of the peripheral stalk, and the F_O_ region. These maps were combined with the map from local refinement of the F_1_ region of the State 1_catalytic_ with αβ_tight_ closed using the ‘vop maximum’ function in *USCF Chimera* to generate a composite map of the entire complex.

### Model building and refinement

To build atomic models of the F_1_ region, the crystal structure of yeast F_1_ (PDB 2HLD)(Kabaleeswaran et al., 2006) was fitted as a rigid body into locally refined maps of F_1_ in *UCSF Chimera* (Pettersen et al., 2004). Models were manually adjusted in *Coot* (Emsley and Cowtan, 2004) before being imported into *ISOLDE* (Croll, 2018) within *ChimeraX* (Goddard et al., 2018) to improve dihedral angles and rotamer fitting. A final round of refinement was performed with *Phenix* and the resulting models were evaluated with *Molprobity* (Chen et al., 2010) and *EMRinger* (Barad et al., 2015). To build backbone models of the full complex, a mosaic model was first assembled by rigid body fitting of the yeast F_1_ crystal structure (PDB 2HLD), subunits abc_10_dfi from the yeast F_O_ cryoEM structure (PDB 6B2Z) (Guo et al., 2017), the peripheral stalk region without subunit h from the yeast monomer cryoEM structure (PDB 6CP3) (Srivastava et al., 2018), and domains of a subunit h atomic model predicted with AlphaFold (Jumper et al., 2021) into the unsharpened maps. Molecular Dynamics Flexible Fitting (Trabuco et al., 2008) was then performed for the 27 rotational states of the dataset including ATP to generate corresponding backbone models. Figures and movies were generated with *ChimeraX* and *UCSF Chimera*.

## Statement of contributions

JLR conceived the project and supervised the research. HG purified the protein, performed the cryoEM, image analysis, and atomic model building. HG and JLR wrote the manuscript and prepared figures.

## Acknowledgements

HG was supported by an Ontario Graduate Scholarship for International Students. JLR was supported by the Canada Research Chairs program. This work was supported by Canadian Institutes of Health Research grant PJT162186 (JLR). CryoEM data was collected at the Toronto High-Resolution High-Throughput cryoEM facility, supported by the Canada Foundation for Innovation and Ontario Research Fund.

## Data availability

CryoEM maps are deposited in the Electron Microscopy Data Bank and atomic models are deposited in the Protein Data Bank with accession numbers indicated in Supplementary Tables 1 and 2.

