## Supporting Information for "Structure of ATP synthase under strain during catalysis"

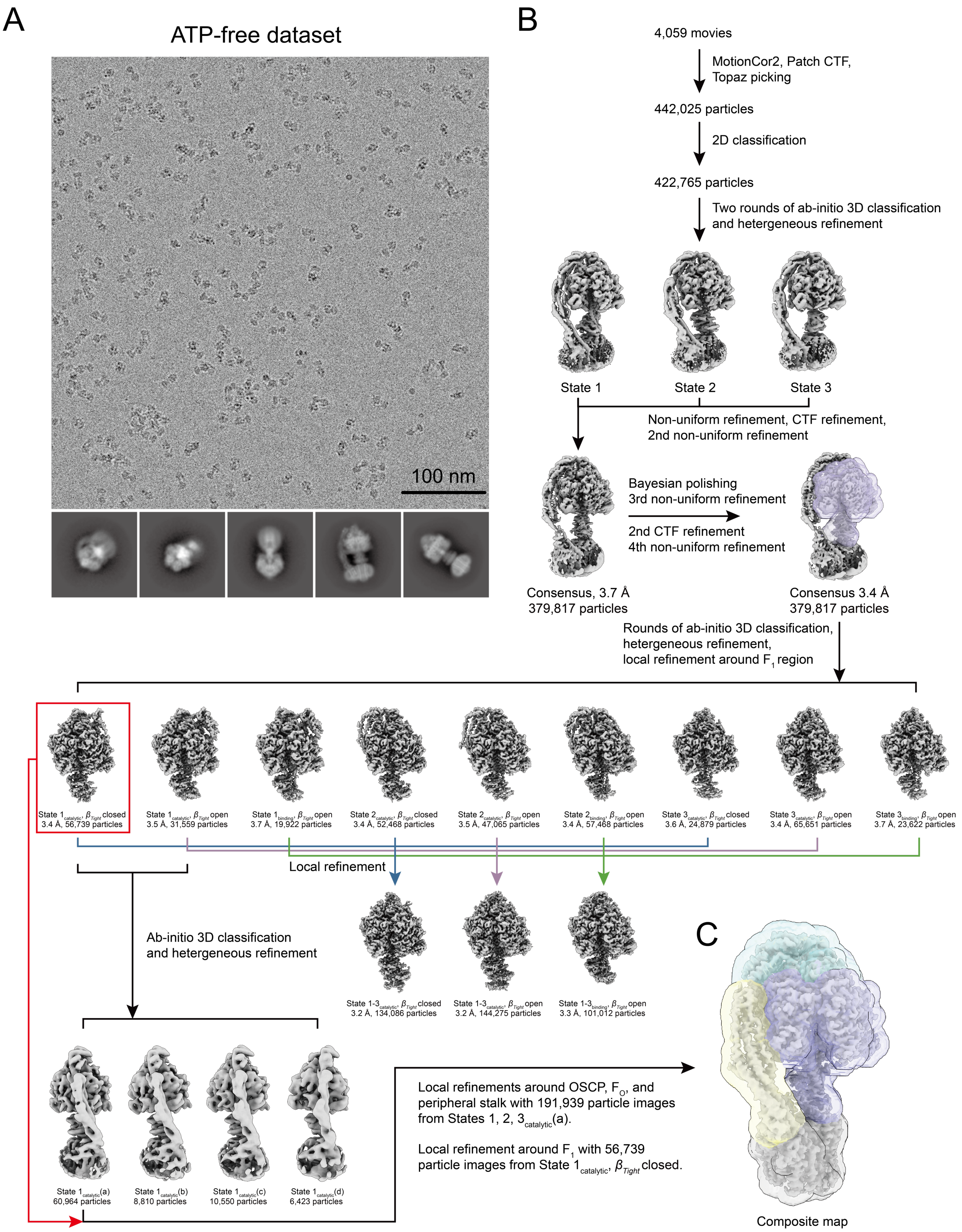


**Supplementary Figure 1. CryoEM of the ATP-free dataset. A,** Example micrograph and class average images. **B,** CryoEM data analysis workflow. **C,** Masks used for focused refinement.

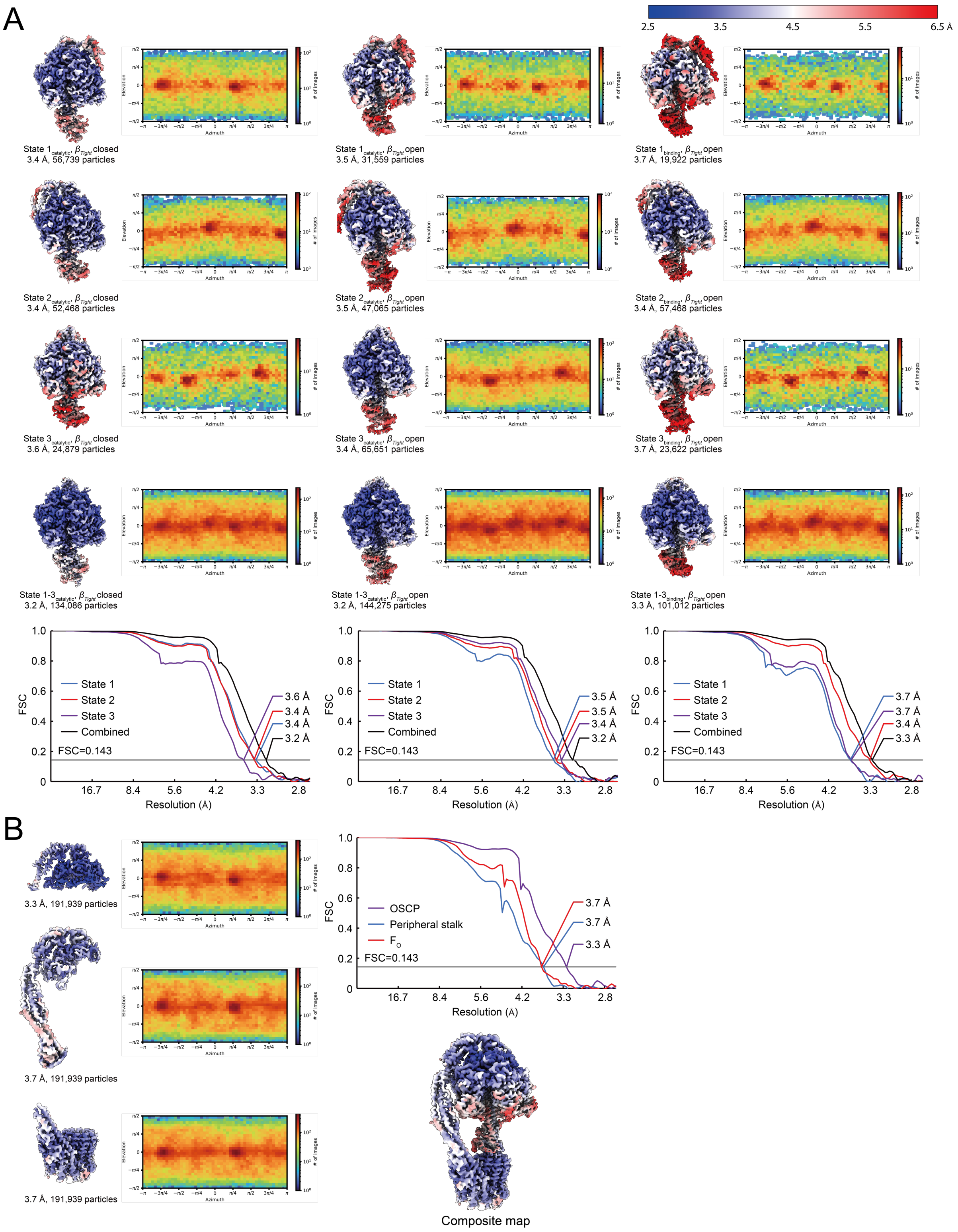


**Supplementary Figure 2. CryoEM map validation for ATP-free dataset. A,** Local resolution maps, orientation distribution plots, and gold standard Fourier Shell Correlation curves (with masking correction) for the different conformations observed. **B,** Local resolution maps, orientation distribution plots, and gold standard Fourier Shell Correlation curves (with masking correction) for focused refinement.

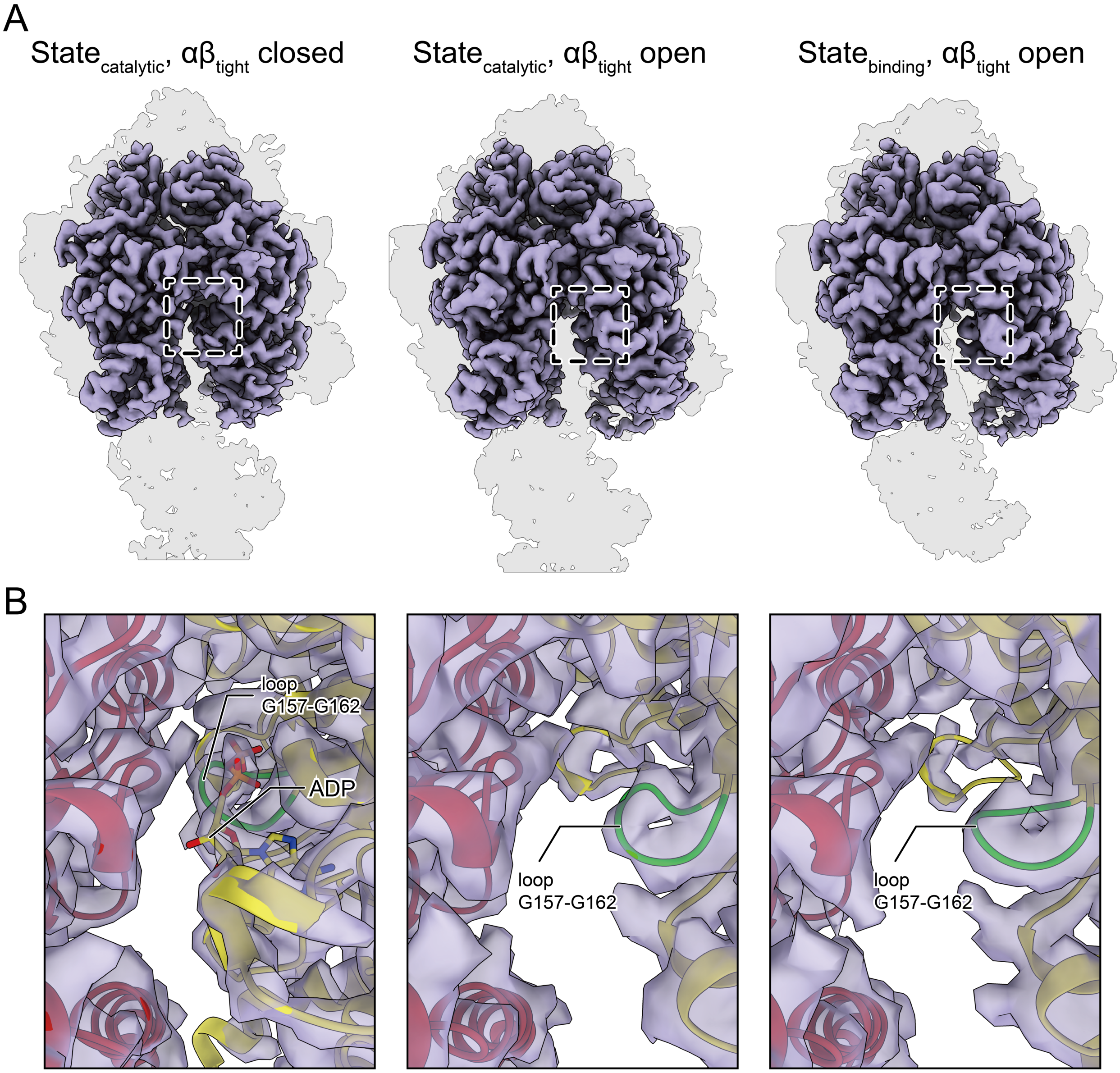


**Supplementary Figure 3: Nucleotide content of the αβ$_{tight}$ site in the absence of free ATP.** **A,** The αβ$_{tight}$ site shows a closed (*left*) or open *(middle*) conformation in the catalytic dwell structure, and an open conformation (*right*) in the binding dwell structure. **B,** Density for nucleotide in the αβ$_{tight}$ site is apparent only for catalytic dwell structure in the closed conformation (*left*).

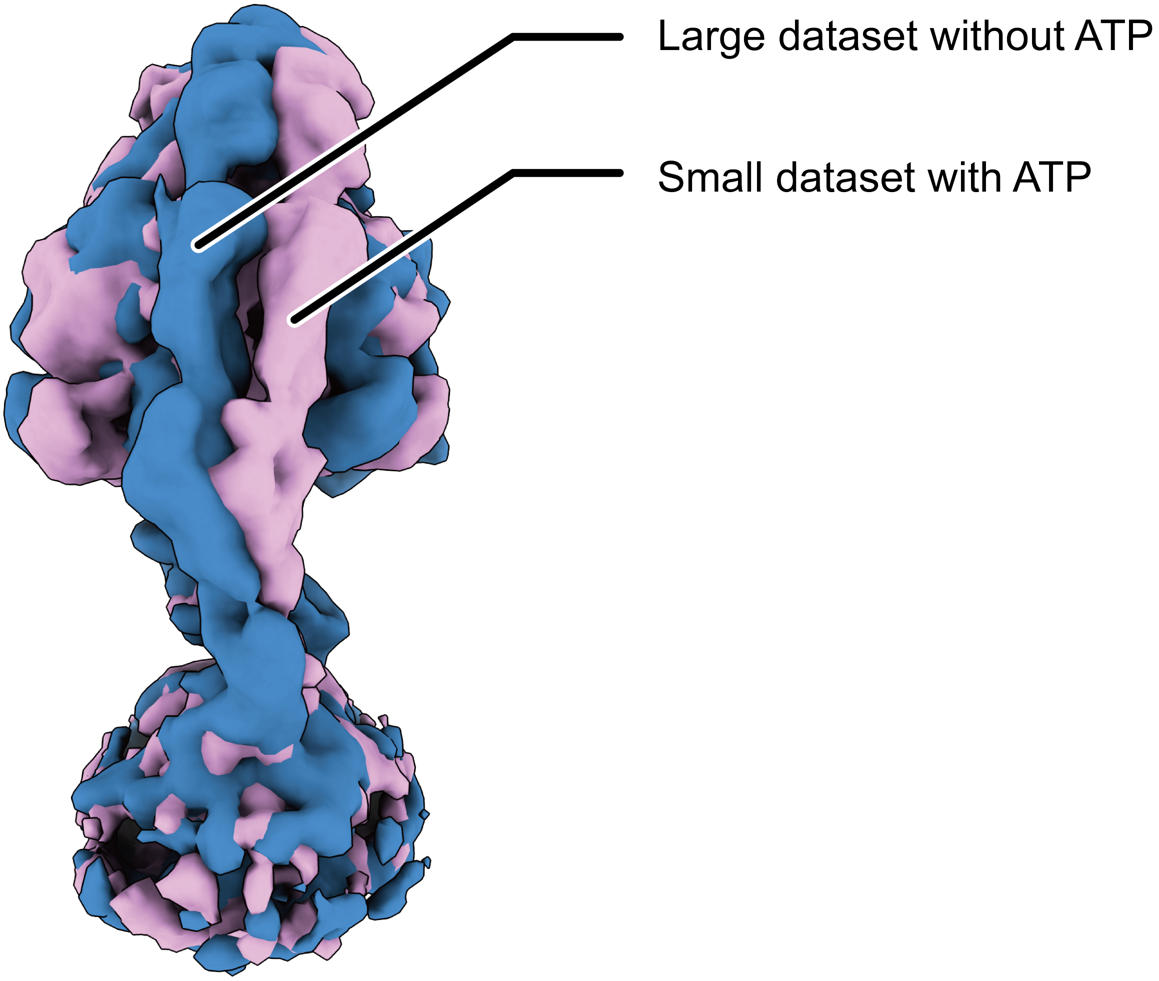


**Supplementary Figure 4. Imaging ATP synthase during ATP hydrolysis reveals conformations with a strained peripheral stalk.** A small dataset of cryoEM images were collected with a screening microscope from ATP synthase frozen during ATP hydrolysis. Analysis of these images yields 3D classes showing the enzyme with a strained peripheral stalk (*pink*), which are not seen in the absence of free ATP (*blue*).

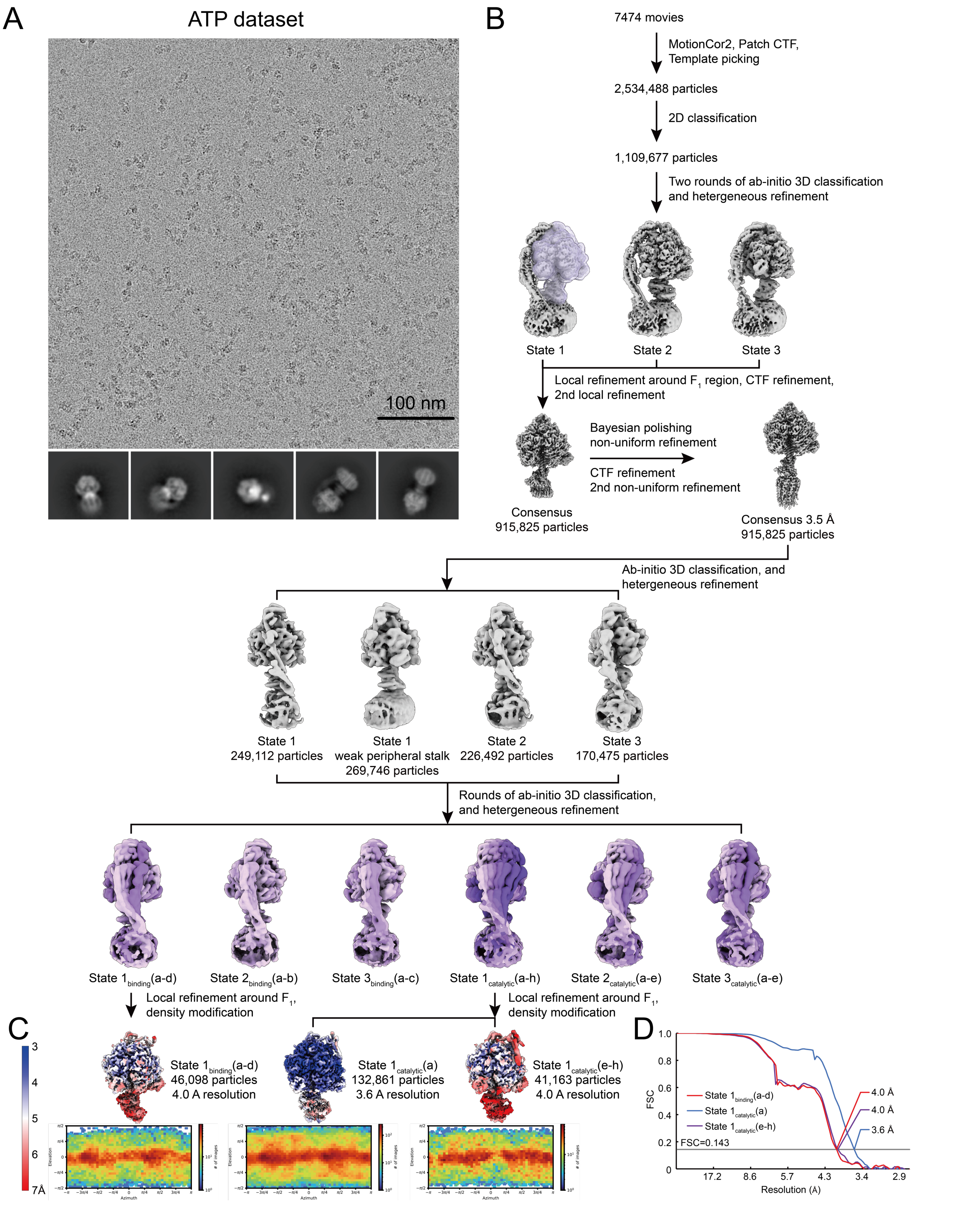


**Supplementary Figure 5. CryoEM of the ATP synthase frozen during ATP hydrolysis. A,** Example micrograph and 2D class average images. **B,** CryoEM data analysis workflow. **C,** Local resolution maps and orientation distribution plots for the $F_1$ region in the different conformations. **D,** Fourier shell correlation curves following gold standard refinement and correction of the effects of masking for focused refinement of the $F_1$ region in the different conformations.

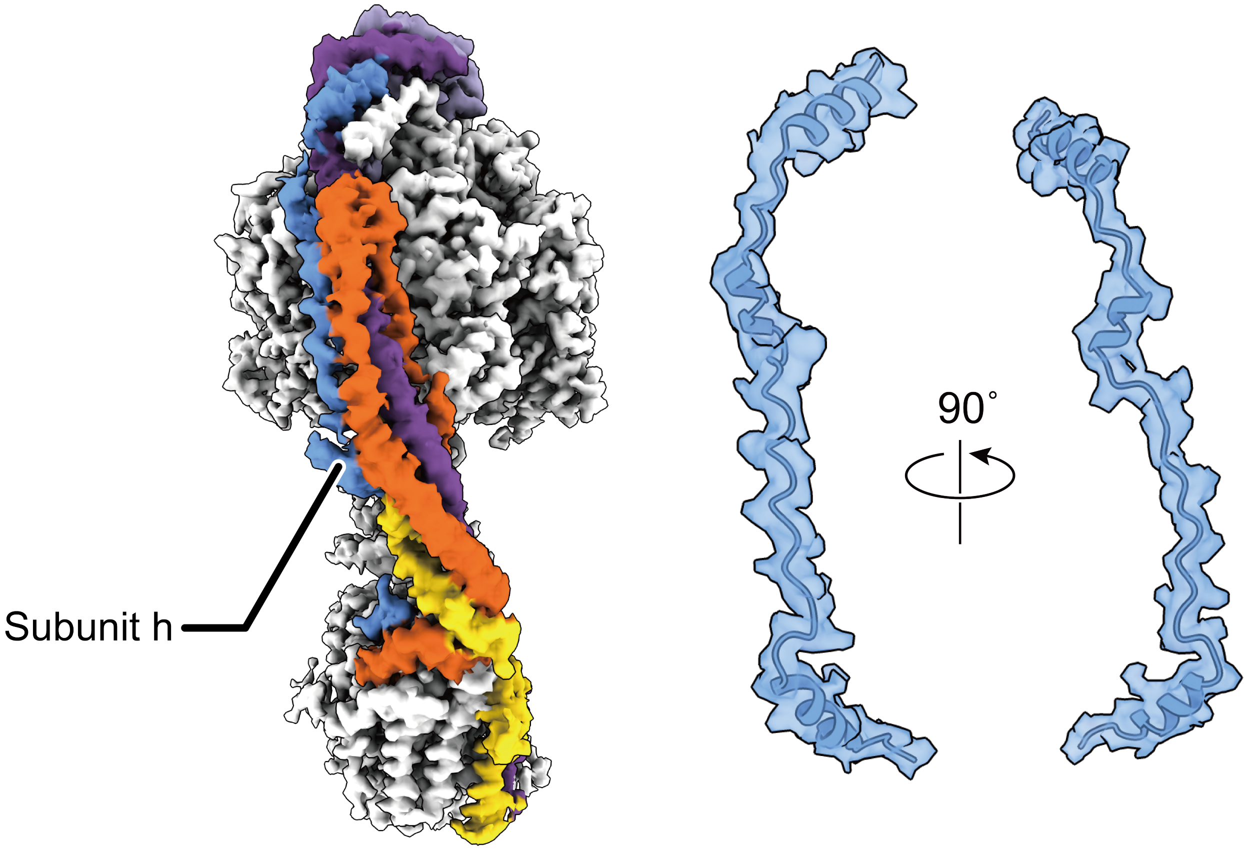


**Supplementary Figure 6. Structure of subunit h in the peripheral stalk.** Focused refinement of the peripheral stalk revealed nearly-continuous density that accommodates a model of residues 1 to 62 of subunit h from AlphaFold.

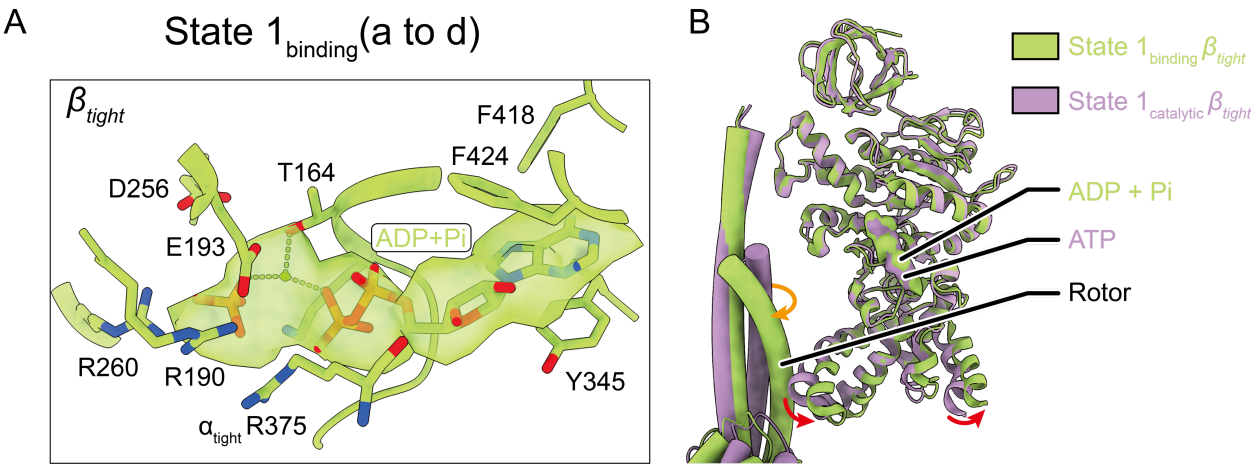


**Supplementary Figure 7. Transition from binding to catalytic states is consistent with a previously proposed model of ATP-hydrolysis induced rotation. A,** Density for nucleotide in the $\beta_{tight}$ site of the binding state is consistent with MgADP and Pi. **B,** The binding and catalytic states different by movement of the β subunit (red arrows) and rotation of the γ subunit (orange arrow).

### Supplementary Table 1. Cryo-EM data collection, refinement and validation statistics

| | ATP free, State 1-3$_{catalytic}$, $\beta_{tight}$ closed | ATP free, State 1-3$_{catalytic}$, $\beta_{tight}$ open | ATP free, State 1-3$_{binding}$, $\beta_{tight}$ open | ATP, State 1$_{catalytic}$(a) | ATP, State 1$_{catalytic}$(e-h) | ATP, State 1$_{binding}$(a-d) |
|---|---|---|---|---|---|---|
| EMDB | 25930 | 25931 | 25932 | 25933 | 25934 | 25939 |
| PDB | 7TJS | 7TJT | 7TJU | 7TJV | 7TJW | 7TJX |
| **Data collection and processing** | | | | | | |
| Magnification | 75,000 | 75,000 | 75,000 | 59,000 | 59,000 | 59,000 |
| Voltage (kV) | 300 | 300 | 300 | 300 | 300 | 300 |
| Electron exposure (e–/Å$^2$) | 39 | 39 | 39 | 40 | 40 | 40 |
| Defocus range (μm) | 0.5-3.4 | 0.5-3.4 | 0.5-3.4 | 0.5-3.0 | 0.5-3.0 | 0.5-3.0 |
| Pixel size (Å) | 1.046 | 1.046 | 1.046 | 1.348 | 1.348 | 1.348 |
| Symmetry imposed | C1 | C1 | C1 | C1 | C1 | C1 |
| Initial particle images (no.) | 442,025 | 442,025 | 442,025 | 2,534,488 | 2,534,488 | 2,534,488 |
| Final particle images (no.) | 134,086 | 144,275 | 101,012 | 132,861 | 41,163 | 46,098 |
| Map resolution (Å) | 3.2 | 3.2 | 3.3 | 3.6 | 4.0 | 4.0 |
| FSC threshold | 0.143 | 0.143 | 0.143 | 0.143 | 0.143 | 0.143 |
| Map resolution range (Å) | 2.9-5.5 | 2.9-5.7 | 2.9-6.1 | 3.0-5.6 | 3.5-7.0 | 3.5-6.6 |
| **Refinement** | | | | | | |
| Initial model used (PDB code) | 2HLD | 2HLD | 2HLD | 2HLD | 2HLD | 2HLD |
| Model resolution (Å) | 3.4 | 3.4 | 3.5 | 3.6 | 3.8 | 3.9 |
| FSC threshold | 0.5 | 0.5 | 0.5 | 0.5 | 0.5 | 0.5 |
| Map sharpening *B* factor (Å$^2$) | -126.0 | -129.5 | -123.3 | -152.1 | -144.0 | -151.3 |
| Model composition | | | | | | |
| Non-hydrogen atoms | 23546 | 23136 | 22696 | 23178 | 22508 | 21671 |
| Protein residues | 3102 | 3099 | 3102 | 3110 | 3097 | 3072 |
| Ligands | 8 | 7 | 7 | 10 | 10 | 11 |
| *B* factors (Å$^2$) | | | | | | |
| Protein | 38.46 | 51.29 | 45.73 | 63.43 | 70.19 | 82.99 |
| Ligand | 38.79 | 46.98 | 41.61 | 59.46 | 61.41 | 75.03 |
| R.m.s. deviations | | | | | | |
| Bond lengths (Å) | 0.003 | 0.003 | 0.002 | 0.003 | 0.003 | 0.003 |
| Bond angles (°) | 0.618 | 0.642 | 0.600 | 0.674 | 0.651 | 0.632 |
| Validation | | | | | | |
| MolProbity score | 0.93 | 1.11 | 0.96 | 0.97 | 1.07 | 1.03 |
| Clashscore | 1.75 | 3.24 | 1.98 | 2.02 | 2.42 | 2.44 |
| EMRinger | 3.14 | 3.19 | 3.19 | 2.69 | 2.06 | 2.15 |
| Poor rotamers (%) | 0.00 | 0.00 | 0.00 | 0.00 | 0.00 | 0.05 |
| Ramachandran plot | | | | | | |
| Favored (%) | 98.54 | 98.83 | 98.57 | 98.06 | 97.79 | 98.19 |
| Allowed (%) | 1.46 | 1.17 | 1.43 | 1.94 | 2.21 | 1.81 |
| Disallowed (%) | 0.00 | 0.00 | 0.00 | 0.00 | 0.00 | 0.00 |

### Supplementary Table 2. Additional maps and backbone models

| | PDB | EMDB | Num. particles | Resolution (Å) | Pixel size (Å) | B factor (Å$^2$) | Details |
|---|---|---|---|---|---|---|---|
| State $1_{catalytic}$, $\beta_{tight}$ closed | | 25935 | 56,739 | 3.4 | 1.308 | -114.0 | Locally refined $F_1$ region generated from the ATP-free dataset (Fig. S1and S2). |
| State $1_{catalytic}$, $\beta_{tight}$ open | | 25936 | 31,559 | 3.5 | ″ | -107.1 | ″ |
| State $1_{binding}$, $\beta_{tight}$ open | | 25937 | 19,922 | 3.7 | ″ | -100.1 | ″ |
| State $2_{catalytic}$, $\beta_{tight}$ closed | | 25938 | 52,468 | 3.4 | ″ | -115.1 | ″ |
| State $2_{catalytic}$, $\beta_{tight}$ open | | 25940 | 47,065 | 3.5 | ″ | -110.9 | ″ |
| State $2_{binding}$, $\beta_{tight}$ open | | 25941 | 57,468 | 3.4 | ″ | -116.2 | ″ |
| State $3_{catalytic}$, $\beta_{tight}$ closed | | 25942 | 24,879 | 3.6 | ″ | -104.6 | ″ |
| State $3_{catalytic}$, $\beta_{tight}$ open | | 25943 | 65,651 | 3.4 | ″ | -113.4 | ″ |
| State $3_{binding}$, $\beta_{tight}$ open | | 25944 | 23,622 | 3.7 | ″ | -99.3 | ″ |
| State $1_{catalytic}$(a) | 7TJY | 25946 | 41,506 | 3.8 | 1.348 | Unsharpened | Substate of State $1_{catalytic}$ generated from the ATP-free dataset. Particle number corresponds to particle images used to calculate the map, not the summed number of particle images in this state (Fig. 1F and S1). |
| State $1_{catalytic}$(b) | 7TJZ | 25947 | 8,810 | 4.4 | ″ | ″ | ″ |
| State $1_{catalytic}$(c) | 7TK0 | 25948 | 10,550 | 4.4 | ″ | ″ | ″ |
| State $1_{catalytic}$(d) | 7TK1 | 25949 | 1,984 | 7.1 | ″ | ″ | ″ |
| OSCP | | 25945 | 191,939 | 3.3 | 1.308 | -128.6 | Locally refined map with particle images from the most relaxed conformations of all three rotational states in the ATP-free dataset (Fig. S1and S2). |
| Peripheral stalk | | 25950 | 191,939 | 3.7 | ″ | -166.3 | ″ |
| $F_O$ | | 25951 | 191,939 | 3.7 | ″ | -173.0 | ″ |
| Composite map | | 25952 | | | ″ | | Composite map of the ATP-free dataset generated with the locally refined maps of the OSCP, peripheral stalk, $F_O$, and the State $1_{catalytic}$, $\beta_{tight}$ closed (Fig. S1and S2). |
| Map from screening microscope | | 25953 | 5,919 | 8.4 | 1.348 | | Map displaying strained peripheral stalk generated with a small dataset collected on the local screening F20 microscope (Fig. S4). |
| State $1_{binding}$(a) | 7TK2 | 25954 | 9,369 | 6.5 | ″ | Unsharpened | Substate generated from the ATP dataset. Particle number corresponds to particle images used to calculate the map, not the summed number of particle images in this state (Fig. 1F and S5). |
| State $1_{binding}$(b) | 7TK3 | 25955 | 8,184 | 6.3 | ″ | ″ | ″ |
| State $1_{binding}$(c) | 7TK4 | 25956 | 6,644 | 7.0 | ″ | ″ | ″ |
| State $1_{binding}$(d) | 7TK5 | 25957 | 3,473 | 7.8 | ″ | ″ | ″ |
| State $1_{catalytic}$(a) | 7TK6 | 25958 | 26,797 | 6.5 | ″ | ″ | ″ |
| State $1_{catalytic}$(b) | 7TK7 | 25959 | 7,581 | 6.7 | ″ | ″ | ″ |
| State $1_{catalytic}$(c) | 7TK8 | 25960 | 16,658 | 4.7 | ″ | ″ | ″ |
| State $1_{catalytic}$(d) | 7TK9 | 25961 | 7,185 | 6.0 | ″ | ″ | ″ |
| State $1_{catalytic}$(e) | 7TKA | 25962 | 9,849 | 7.1 | ″ | ″ | ″ |
| State $1_{catalytic}$(f) | 7TKB | 25963 | 6,516 | 6.3 | ″ | ″ | ″ |
| State $1_{catalytic}$(g) | 7TKC | 25964 | 8,727 | 5.8 | ″ | ″ | ″ |
| State $1_{catalytic}$(h) | 7TKD | 25965 | 3,251 | 7.7 | ″ | ″ | ″ |
| State $2_{binding}$(a) | 7TKE | 25966 | 14,051 | 7.1 | ″ | ″ | ″ |
| State $2_{binding}$(b) | 7TKF | 25967 | 9,157 | 7.1 | ″ | ″ | ″ |
| State $2_{catalytic}$(a) | 7TKG | 25968 | 32,285 | 4.5 | ″ | ″ | ″ |
| State $2_{catalytic}$(b) | 7TKH | 25969 | 25,401 | 4.4 | ″ | ″ | ″ |
| State $2_{catalytic}$(c) | 7TKI | 25970 | 4,213 | 7.1 | ″ | ″ | ″ |
| State $2_{catalytic}$(d) | 7TKJ | 25971 | 3,208 | 7.5 | ″ | ″ | ″ |
| State $2_{catalytic}$(e) | 7TKK | 25972 | 3,761 | 7.3 | ″ | ″ | ″ |
| State $3_{binding}$(a) | 7TKL | 25973 | 12,214 | 6.4 | ″ | ″ | ″ |
| State $3_{binding}$(b) | 7TKM | 25974 | 21,305 | 4.5 | ″ | ″ | ″ |

| | | | | | | | |
|---|---|---|---|---|---|---|---|
| State $3_{binding}$(c) | 7TKN | 25975 | 16,301 | 7.1 | ″ | ″ | ″ |
| State $3_{catalytic}$(a) | 7TKO | 25976 | 11,541 | 4.8 | ″ | ″ | ″ |
| State $3_{catalytic}$(b) | 7TKP | 25977 | 16,774 | 4.6 | ″ | ″ | ″ |
| State $3_{catalytic}$(c) | 7TKQ | 25978 | 24,660 | 4.5 | ″ | ″ | ″ |
| State $3_{catalytic}$(d) | 7TKR | 25979 | 10,914 | 6.5 | ″ | ″ | ″ |
| State $3_{catalytic}$(e) | 7TKS | 25980 | 3,435 | 7.5 | ″ | ″ | ″ |
